# A hidden network of gene relationships unifies behavioral and molecular research

**DOI:** 10.1101/2025.11.14.688547

**Authors:** David R. Connell, Chris Gaiteri, Shinya Tasaki

## Abstract

Behavioral and molecular approaches often share topics of inquiry across basic and applied research fields. However, for historical and practical reasons, these approaches conceptualize scientific questions differently. This segregation is particularly damaging to translational fields that are in need of the combined strength of these two major approaches. While studies are increasingly recognizing the value of integrated approaches through hybrid methods, these efforts require great resources. To potentially relieve this interdisciplinary detachment, we uncover similarities between behavioral and molecular work in the publication network to test if the two approaches are more aligned than they appear, through deeper common concepts. To do this, we first create a model to predict genes that are relevant to behavioral publications from their abstracts. These predictions are in turn used to define gene-based similarity across the entire medical literature network. This model is built on-top of a Large Language Model (LLM), used to find patterns in the semantic structure of abstracts. Our experiments demonstrated the model’s efficacy in predicting gene annotations in molecular abstracts and showed the practical value of predictions made on behavioral research. We found similar increased relative risks for differentially expressed genes in disease-specific publications compared to the global publication population in both the molecular and behavioral groups. Additionally, the model was able to correlate gene predictions between molecular publications and their behavioral references. We instantiated this model in a free and easy to use search engine that finds related publications via the latent representations we’ve established. Using this model, Abstract2Gene facilitates novel collaborations by uncovering shared gene signatures across medical literature, broadening the reach of research to distant fields and promoting collaboration between molecular and behavioral researchers.

## Introduction

There is resistance to information flow between molecular and behavioral researchers (Table 1) despite related topics of study. Disease symptoms at a clinical level have been observed to predict their molecular level roots [1]. More generally, systems biology has repeatedly produced strong evidence relating genes, proteins, diseases, and symptoms highlighting the importance of their relationships [1–4]. This indicates the value of the multi-scale network tying molecular physiology to behavioral level consequences as opposed to viewing each in isolation. The preference for within field research seen in the publication network limits discovery and generating work capable of disrupting current thinking [5, 6]. To maximize the efficiency of medical research, the distant points of the publication network need to be more available to promote cross-discipline collaboration. Aiming to address this challenge, we have created a model to identify relationships between behavioral and molecular research by predicting previously hidden gene associations in publications to mirror the biological multi-scale networks in the publication landscape.

**Table 1.** Distribution of references in the PubMed dataset. The percentages of references from a publication type (behavioral or molecular) to their references’ types are presented next to the raw counts. This dataset contains data on 24.6 million publications with reference data for 9 million publications.

| Publication type | Molecular references | Behavioral references | Total references |
| --- | --- | --- | --- |
| Molecular | 75,468,816 (66%) | 38,727,825 (34%) | 114,196,641 (100%) |
| Behavioral | 37,513,924 (22%) | 133,831,783 (78%) | 171,345,707 (100%) |
| All | 112,982,740 (40%) | 172,559,608 (60%) | 285,542,348 (100%) |

Gene annotations have been created for the publications in the PubMed dataset in the past including ”gene2pubmed”, an attempt at manually labeling publications, and ”gene2pubtator3”, which is aided by a Name Entity Recognition (NER) model to automate the arduous task [7]. Both these annotation sets have been invaluable for text based analyses of the medical literature, but they both label based on a publication’s direct association with a gene. Additionally, all gene annotations are treated equivalently, labels are binary instead of weighing the importance of a gene to a publication. We hypothesize that medical research which does not directly study genes still contains latent gene components based on the relationship between symptoms and genes. To test this hypothesis, we created an LLM based model that uses patterns in abstracts to predict genes.

This model was used to predict genes for all publications in the PubMed dataset. We designed a series of experiments to test if the predictions for genes associated with behavioral publications (here those that were not given a gene annotation by PubTator3) have practical value. First molecular publication predictions were compared to PubTator3’s annotations to show the gene predictions were not random.

To validate model’s predictions on behavioral publications, we compare the gene predictions given to molecular publications with the predictions of their references. We found the vector of gene predictions for both molecular and behavioral references have higher correlation with their referencing publication than randomly selected publications. Further testing the citation network’s relationships, we looked at the gene with the highest prediction in molecular publications and compared that to the prediction of the same gene in their references, this showed, in both molecular and behavioral publications, the prediction of the gene was correlated with the prediction in the referencing publication.

In the last experiment, we further show the model is able to generalize out to behavioral publications and show the functional value of the gene annotations in a real-world application. We used gene annotations from molecular publications to identify genes that show differential expression of transcripts in the Alzheimer’s Disease (AD) population which are also at an elevated Relative Risk (RR) to be found in publications on AD compared to the general publication population. These genes are then compared to genes with a high RR based on the predicted annotations given to behavioral publications. We find the overlap between the genes identified by the molecular publication network and those found by the behavioral publication network. Additionally, the models are able to detect different and more Differentially Expressed (DE) genes than the original PubTator3 annotations.

These experiments give strong evidence the model is able to make meaningful predictions of genes related to behavioral publications. With confidence in the quality of the predictions, we used the vectors of gene predictions as the bases of similarity for a search engine. Our search engine accepts a paper’s title and abstract, allowing a researcher to pass in their working abstract, and the model is used to extract information about which genes it is related to which it uses to suggest potential references across domains based on indirect overlap of genes. This is intended to mimic the networks found in the human disease network to recommend publications in a way not possible with tradition search engines.

## Results

### Creating a model to predict genes in abstracts

To develop a search engine that finds references based on common gene components, we needed to be able to predict genes in abstracts that can be used to calculate similarity across the publication network. We designed a model by combining an LLM with a single linear layer of 768 features. We chose the LLM after training 7 pre-trained LLMs to distinguish abstracts based on whether they are associated with a common PubTator3 gene annotation or not. After training, we chose our fine-tune of the Specter2 model, which was initially trained entirely on scientific literature from the Semantic Scholar database [8].

The Abstract2Gene (A2G) model generates a 768-dimension vector for each abstract. Predictions for multiple genes were calculated from this single vector by comparing the vector to gene-specific templates. We created these templates by averaging the embeddings from ”template size” abstracts sharing a common PubTator3 gene annotation. By averaging many abstracts, we intended on removing the document specific details while emphasizing common gene-related features. To make predictions, we correlated the test abstract’s vector against each gene-specific template and applied a sigmoid function to the results to produce a prediction between 0 and 1 for each gene (Fig 1).

**Figure 1.**
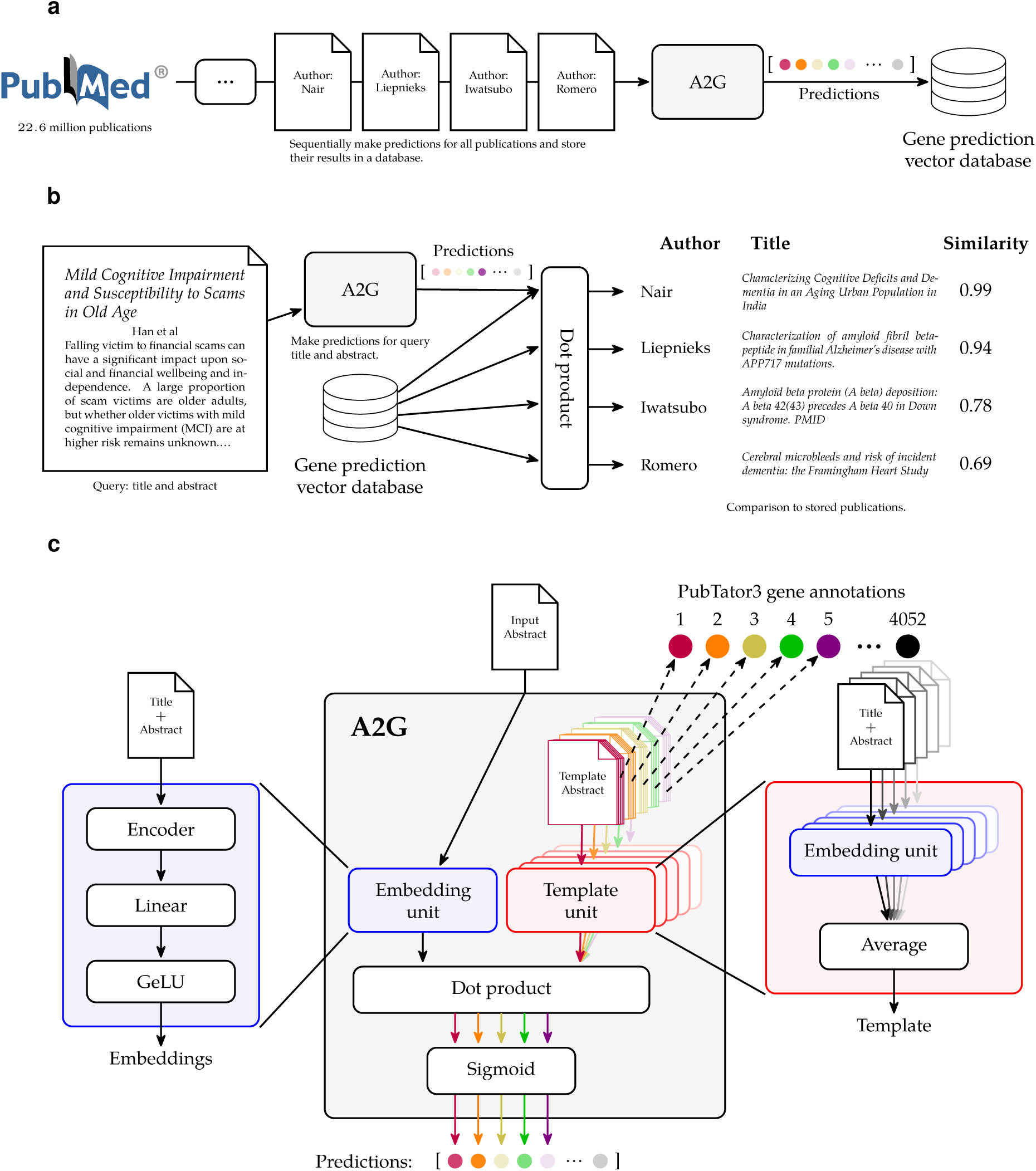
Abstract2Gene model architecture. A. Building the database: All million abstracts from the PubMed dataset were passed to the model, serially, and the predictions were stored in a vector database to allow for quick comparisons. B. Discovering references: A title and abstract is passed to A2G, which predicts the genes related to the input and compares it to all the pre-calculated predictions in the vector database. C. Abstract2Gene model: The A2G model expects a title and abstract as an input. It passes the input document through the embedding unit and compares to each gene template to make a prediction for that gene. The templates are created for each gene by passing several publications associated with that PubTator3 gene annotation through the embedding unit and taking the average embedding.

The dataset we used during this training included 655,552 molecular publications with 18,638 unique genes after converting genes to human orthologs. We dropped genes that were found in less than 8 publications since they don’t have enough examples to create templates, leaving 9124 unique genes. We found the number of a gene’s occurrences in the full dataset to span an exponential range (Fig S1) with most genes mentioned in 10–1000 publications but the most frequent genes have upwards 1 × 10^5^ annotations. In the training dataset, the most common gene was tumor necrosis factor (*TNF*) having 28,362 annotations. Because the number of individual genes was on the order of 1000, label vectors were sparse, which posed a challenge. When all labels were used during training, the zero dominated label vectors biased the model toward low predictions (predicting all zeros achieves an accuracy near 100%). To avoid this, we trained the model on batches with only a small number of labels. We split the labels evenly across the batch size—if there was a batch size of 128 and 8 labels per batch, each label in the batch would get 16 samples. To test the impact of labels per batch, we trained models using different powers of 2, from 2–256 labels per batch and compared the results.

The A2G model trained using 2 labels per batch made predictions that aligned well with the PubTator3 annotations but scores for matching and non-matching publications had high variance (Fig 2). While the sample means for predictions on non-matching publications was consistently lower than the sample means of matching publications for a given gene, the samples could not be perfectly discriminated by a threshold. Additionally, some genes had a stronger signal that the model was able to pick out than others. Using *JAK2* as an example, the model was consistently able to predict negatives near 0 while most of the positives were given predictions above 0.5 but there were a few positive predictions much closer to 0 than the others. These low scoring positives may be cases where PubTator3’s goal of detecting mentions of gene symbols in an abstract do not match with our goal of determining the importance of a gene in the publication. For example, the lowest scoring abstract (PubMed IDentifier (PMID): 31896782, score: 0.014) focuses on the acute myeloid luekemia and the core binding factor (CBF) factor but identifies gene mutations (including mutations of *JAK2*) that may cooperate with previous leukemia founder mutations to advance leukemia progression. While PubTator3 does correctly identify the *JAK2* symbol, this is a secondary gene in the publication and we want our model to give it a lower score to reflect this. In contrast, the model predicts the primary genes, *CBFB* and *RUNX1*, with scores of 1.000 and 0.992 respectively. Additionally, in the highest scoring non-matching abstract (PMID: 21258401, score: 0.670), *STAT3*, a gene closely related to *JAK2* in the JAK2/STAT3 signaling pathway, activation is a key aspect of the findings.

**Figure 2.**
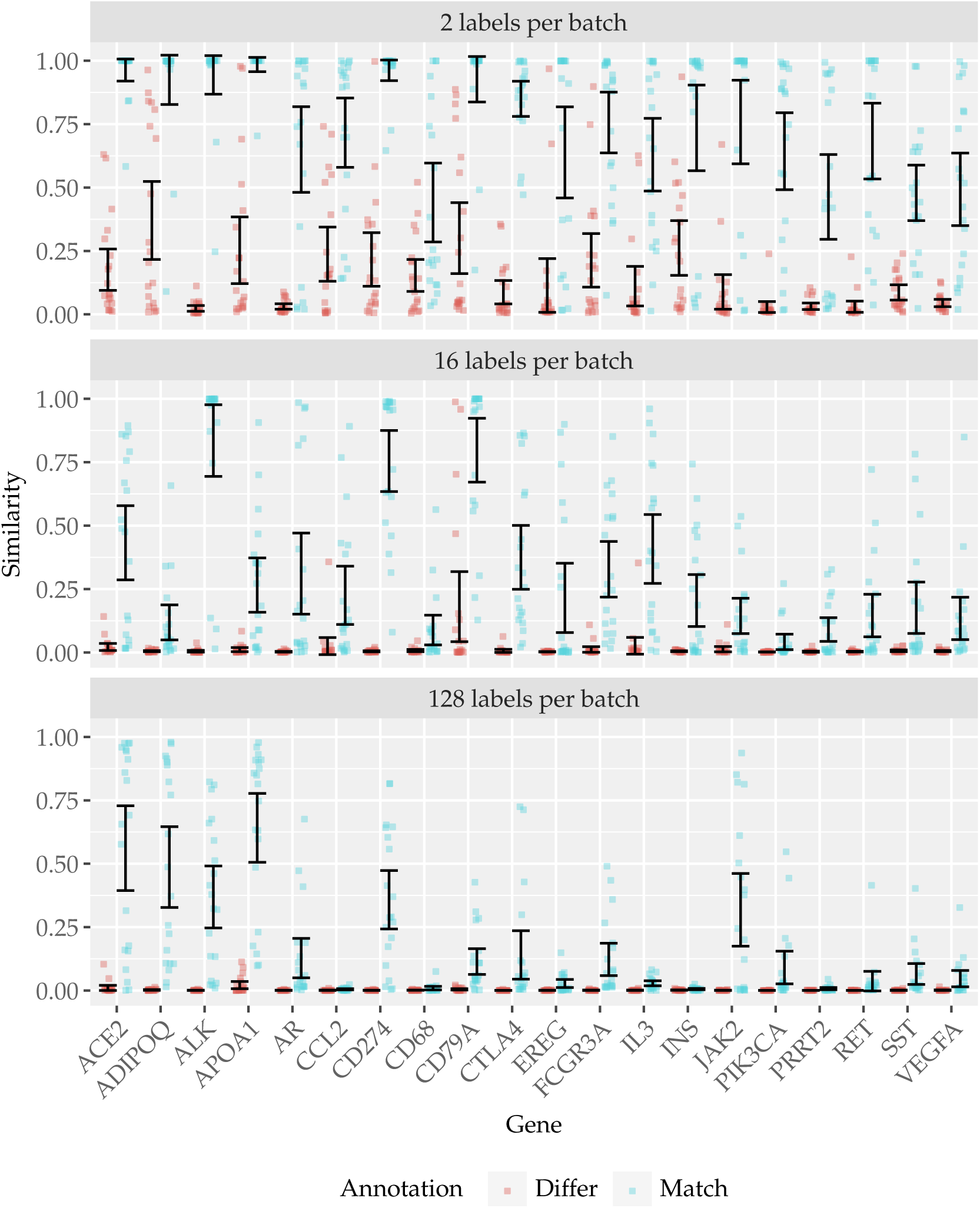
Predicting PubTator3annotations. Performance of A2G trained with 2 (top), 16 (middle), and 128 (bottom) labels per batch against PubTator3’s annotations. For a random sample of 18 genes, 30 examples of publications annotated with that gene (match) were picked and 30 examples without that annotation (differ) were predicted by the model. The error bars show the 95% confidence interval for the mean.

For another gene, *VEGFA*, negatives were given low predictions but there was a wide spread of predictions given to positive examples, one explanation could be that *VEGFA* is often a supporting gene in publications, frequently studied alongside other genes that are more central to the publication, giving *VEGFA* varying degrees of influence on the overall embedding. We looked at the middle scoring matching-abstracts for this gene to better understand the cause of this wide-range of scores. In PMID: 34671630 (score: 0.539) and PMID: 21445393 (score: 0.517) *VEGFA* is one of a list of factors studied on an effect, justifying a lower score suggesting some influence on the publication. For publications PMID: 9405464 (score: 0.573), PMID: 37247185 (score: 0.424), and PMID: 30497276 (score: 0.371), however, *VEGFA* and its receptor are more central. A second reason for a large range of predictions for genes with a PubTator3 annotation associated with the abstract could be their involvement in varied research topics. A gene studied in terms of a distinct single function should be easier to predict than one that is involved in many pathways.

When we increased the labels per batch to 16, the prediction values decreased globally. This was particularly pronounced in the negative case where negative examples’ predictions were increasingly squeezed towards 0. With 16 labels per batch, false positives were rare at the cost of more false negatives. This indicates the model has more confidence when giving the high scoring negatives for *CD79A* suggesting the abstracts do have a relationship with the gene. When comparing 2 and 16 labels per batch, 16 labels per batch emphasized genes with clearer influence on the embeddings (*ALK* and *CD274*). Further increasing the labels per batch to 128 produced a model that prioritized confidence, giving high scores for genes with strong signals while ignoring noisier signals. In this sense, similarity calculated by models with lower labels per batch could be thought of as more creative but less reliable than those calculated with a higher label per batch model. Notably, with 128, the non-matching abstracts that scored high with the 16 label per batch model were clustered close to 0 along with the other non-matching abstracts while some of the matching abstracts were still given scores higher than the non-matching group.

The comparison of models trained on different labels per batch shown in Fig 3, can be used as one factor in deciding the model to use for the search engine. When selecting a model, we balanced false positive rates (acknowledging potential PubTator3 annotation gaps) against maintaining high scores for true positives. These criteria suggest a model between 8 and 64 labels per batch. However, data capping due to high predictions can degrade the performance of the model. We removed 8 labels per batch due to its large number of scores near one.

**Figure 3.**
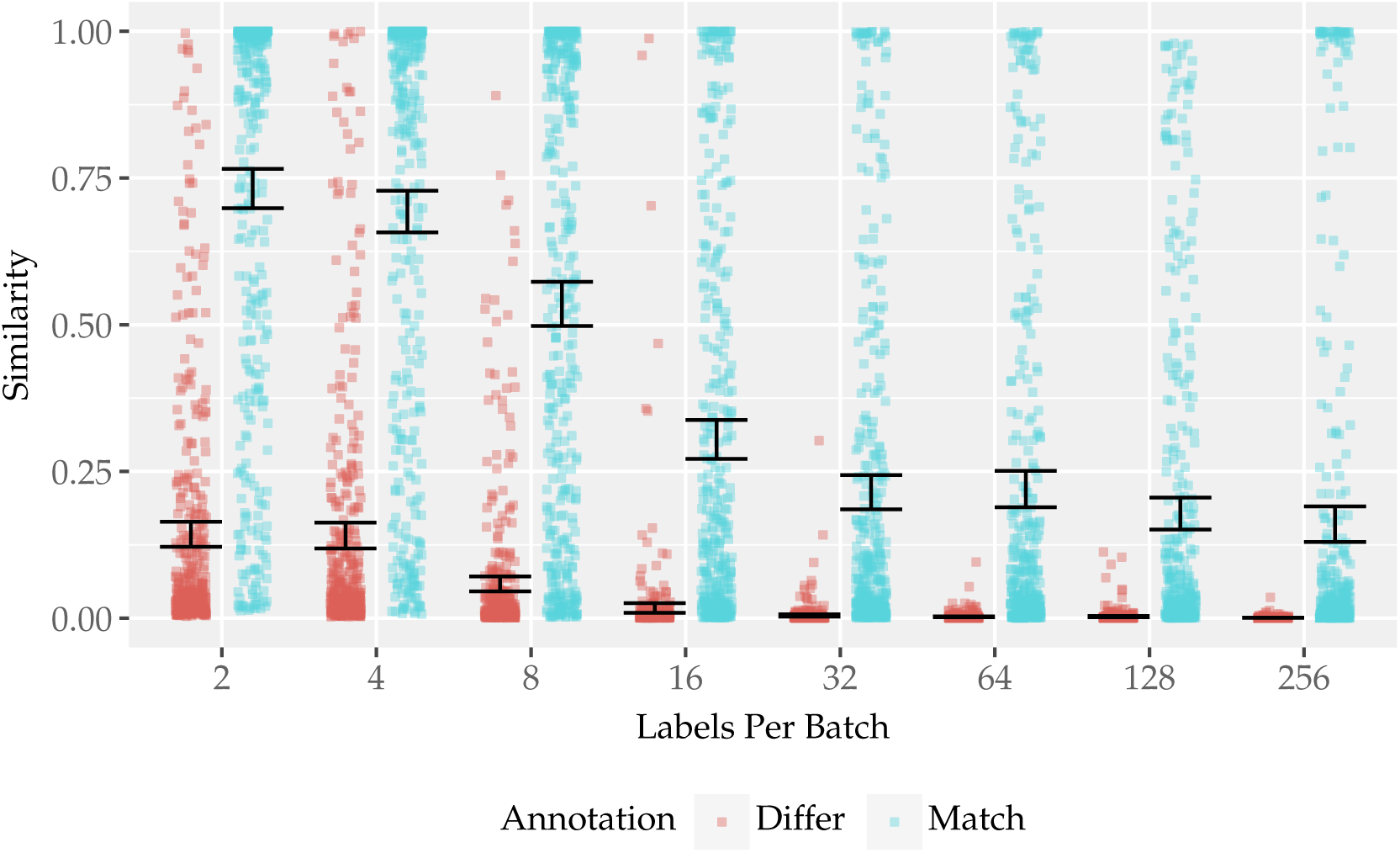
Comparison of all predictions across models trained with different labels per batch. Match samples are predictions for a gene the publication has a PubTator3 annotation for while differ samples are prediction for a publication without a PubTator3 annotation for that gene. All publications used have at least one gene annotation.

### Relating molecular gene predictions to behavioral references

Without gene annotations for behavioral references, we developed an indirect method to validate our model’s ability to make valuable predictions. We hypothesized that molecular publications would cite papers that share gene components, so we leveraged the citation network to compare gene vectors between publications and their references. This approach demonstrated that our model’s predictions matched papers and their citations and that behavioral publications did receive biologically relevant gene scores.

We performed this experiment by randomly selecting molecular publications as ”parent publication” (the citing papers). From each parent reference list we then pulled one molecular and one behavioral reference, as well as a random behavioral publication not referenced by the parent. We then compared each parent to all three child publications using Pearson correlation of their gene vectors.

Using 2 labels per batch to train the model, Fig 4 shows a strong positive shift in similarity when the behavioral publication was referenced by the parent publication compared to a random behavioral publication. Unexpectedly, some random behavioral publications had high similarity with the parent, with some exceeding 0.75. This is likely a consequence of the low label per batch model which favors high predictions across all genes. As anticipated, molecular references had a stronger correlation with the parent than behavioral references, due to their greater focus on genes.

**Figure 4.**
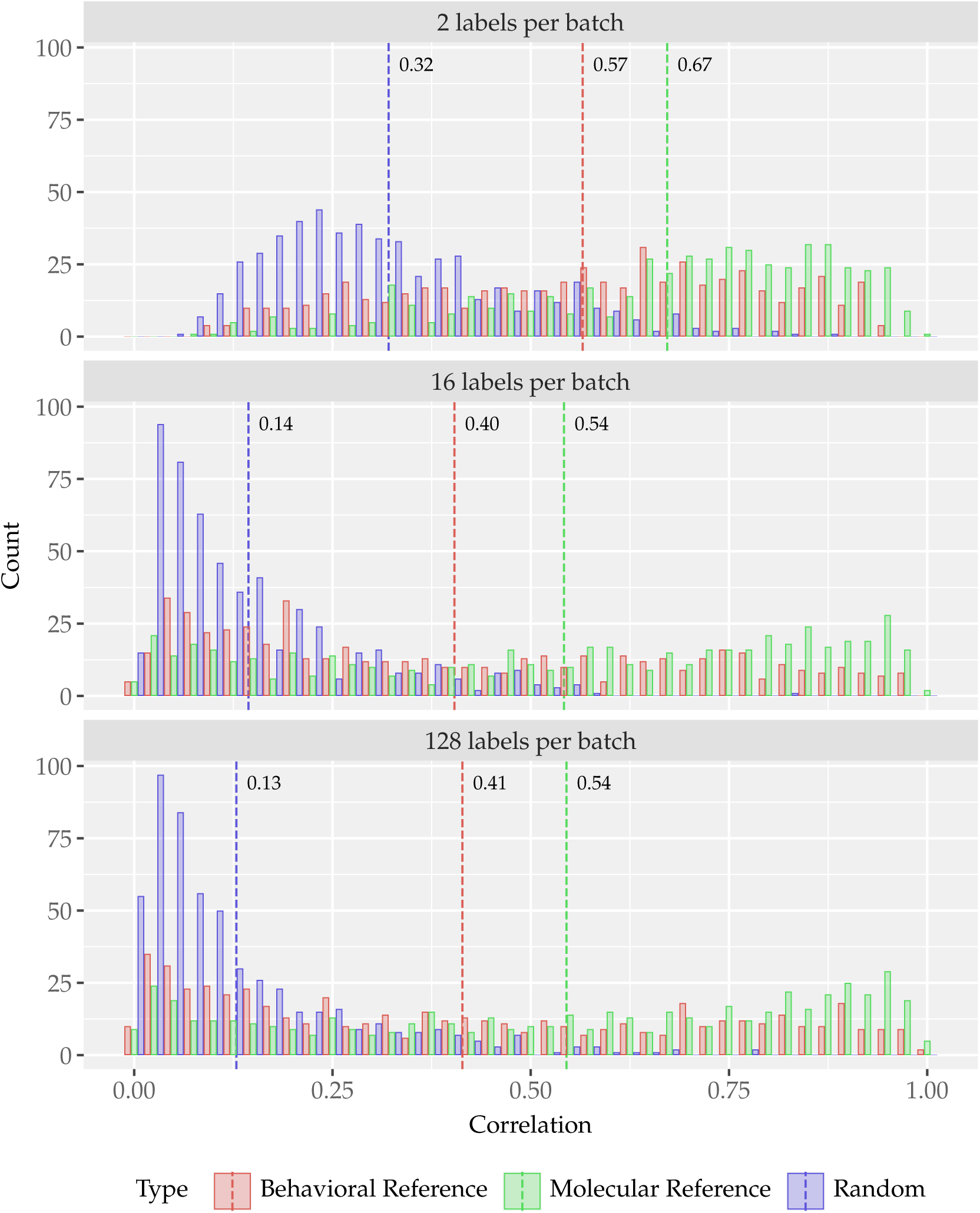
Correlation distribution of molecular parent publications. Using A2G trained with 2 (top), 16 (middle), and 128 (bottom) labels per batch to predict gene vectors. Each sampled parent publication is compared to one of it’s molecular references, one behavioral reference, and another behavioral publication not referenced by the parent. Dotted lines show the mean correlations.

When we trained models with higher labels per batch, the distribution of the random publications’ similarities dropped and became more concentrated. With 128 labels per batch Fig 4 bottom, few random publications have a correlation > 0.5. When comparing models, Fig 5 we observed models trained on 16–128 labels per batch have similar distributions. Interestingly, the pattern as labels per batch increases was not consistent. A2G trained on 4 labels per batch has higher correlation than 2 labels per batch on all groups and the random group is particularly increased compared to 2 labels per batch.

**Figure 5.**
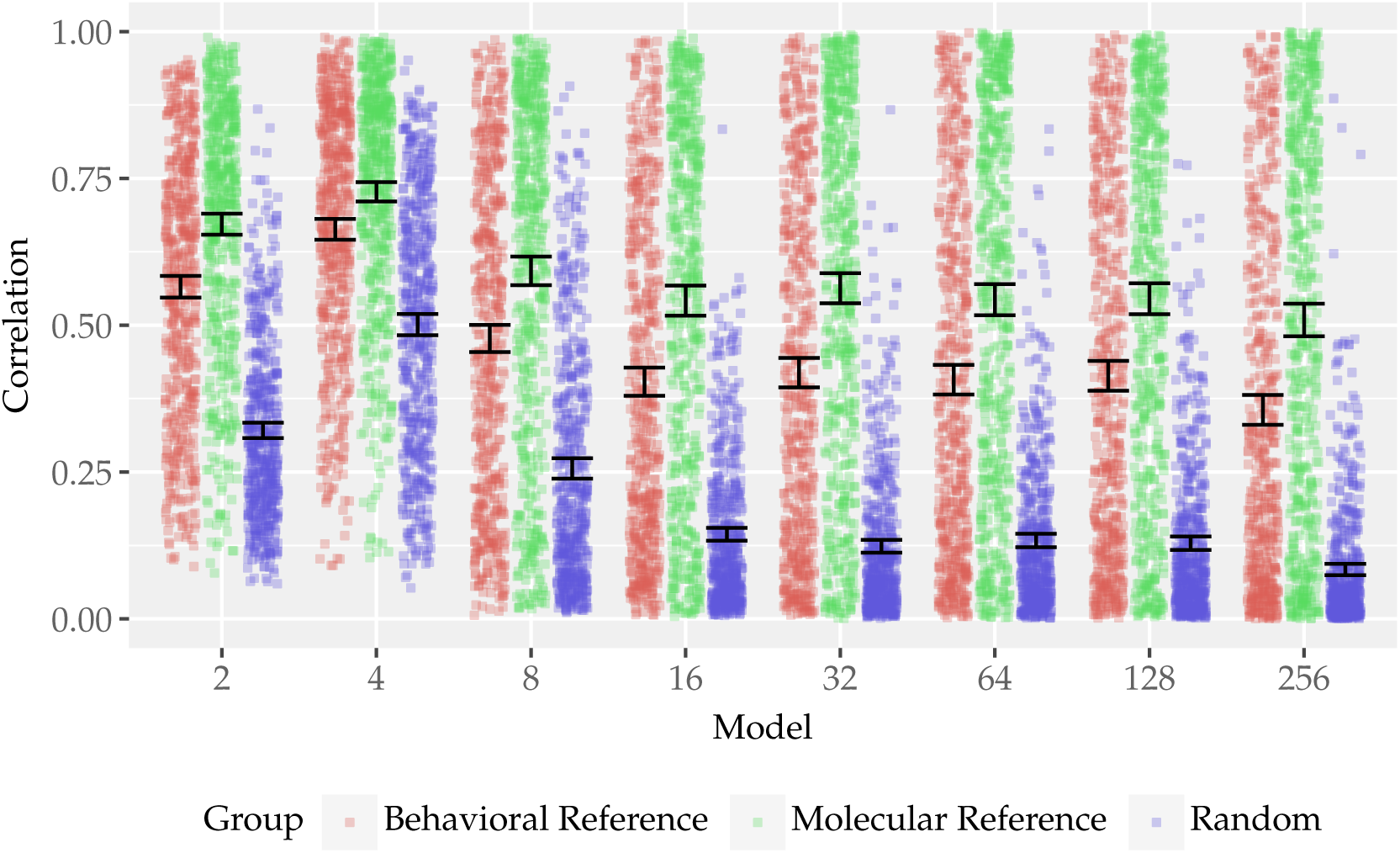
Correlation distribution of models trained with different labels per batch. Comparison of the correlation distributions of models trained with ^1…9^ labels per batch.

To examine gene-specific patterns, we analyzed the top-scoring gene in the parent publications and correlated it with the same scores in the child publications (Fig 6). There was a clear pattern as the parent’s top gene prediction increased, the prediction of the same gene in the references also increased. While molecular references tended to have a greater correlation with their parent publication, we found the same trend of a reference’s gene score increasing as the parent’s score increased in both molecular and behavioral publications.

**Figure 6.**
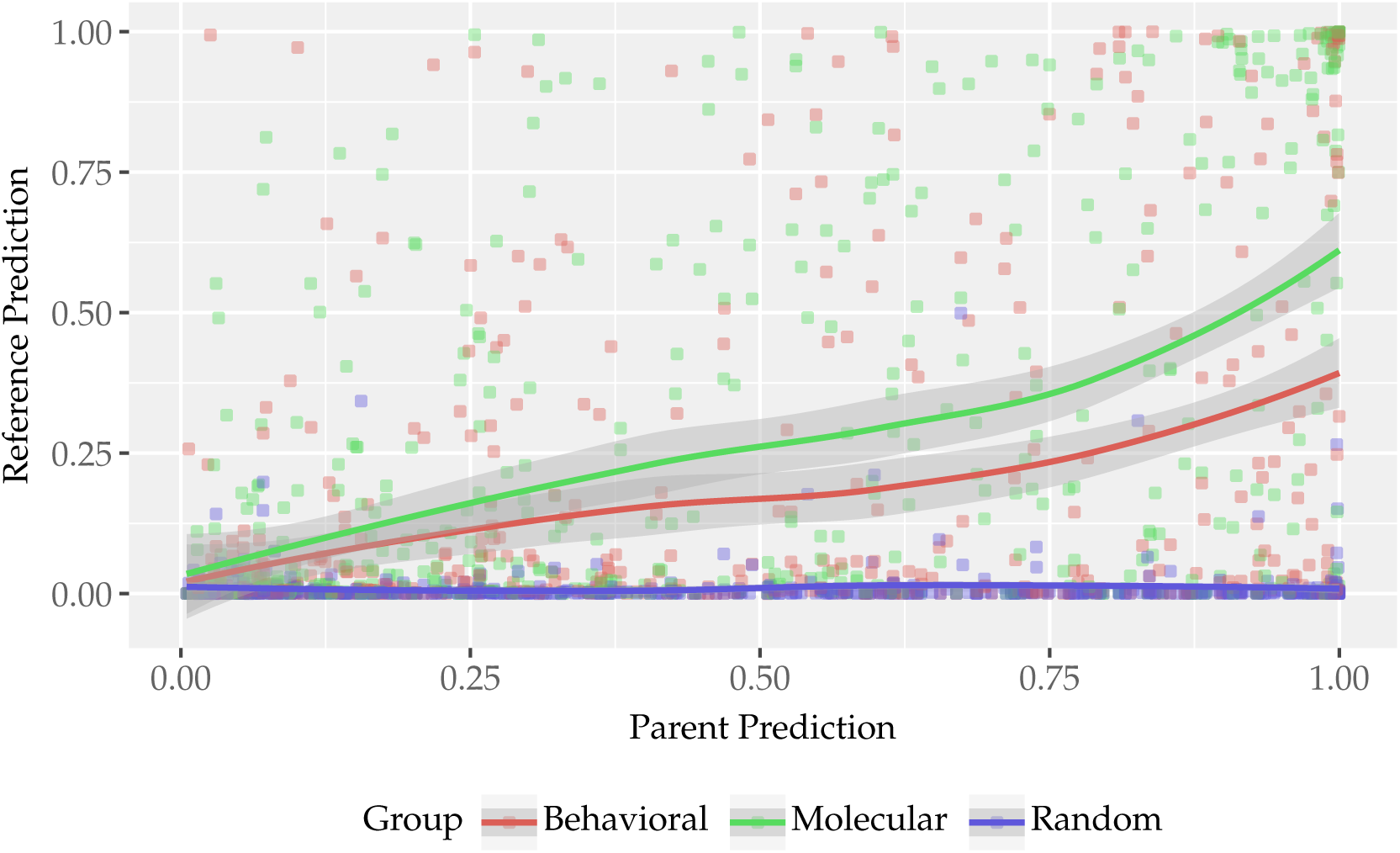
Correlation between top scoring genes. Correlation between the parent reference’s top scoring gene and the prediction for the same gene in the child publications. Predictions performed with a model trained using 128 labels per batch.

### Evaluating molecular vs behavioral Alzheimer’s disease differential gene expression predictions

To evaluate the practical value of the model’s predictions, we created an experiment to test the model’s ability to identify DE genes using molecular and behavioral publications. First, we estimated genes with DE in the medial frontal cortex between participants with AD and Non-Clinically Impaired (NCI) participants using data from the ROSMAP studies [9]. We then used each A2G model to predict the genes related to AD and non-AD publications and tested whether the likelihood a publication was associated with AD DE genes was higher in AD related publications compared to the general population of publications. To perform a RR analysis on the publication gene predictions, we binarized the gene vector with thresholds calculated to match the total number of PubTator3 annotations. To determine whether the model works on behavioral publication, we performed this test twice, once on only molecular publications and again on only behavioral publications and compared the results.

Our first analysis used only molecular publications and compared the binarized A2G predictions to PubTator3’s gene annotations (Fig 7 top). The *MAPT* gene was the only gene all annotation methods agreed had a RR > 1 for AD publications. High RR genes were not consistent across A2G models but there was overlap in several genes including *ADAM10*, *PNPLA6*, and *TOMM40*. Given the 296 DE genes identified, this was unlikely to be due to random chance. The A2G models trained on 64 and 256 labels per batch found the most high RR genes at 6 each.

**Figure 7.**
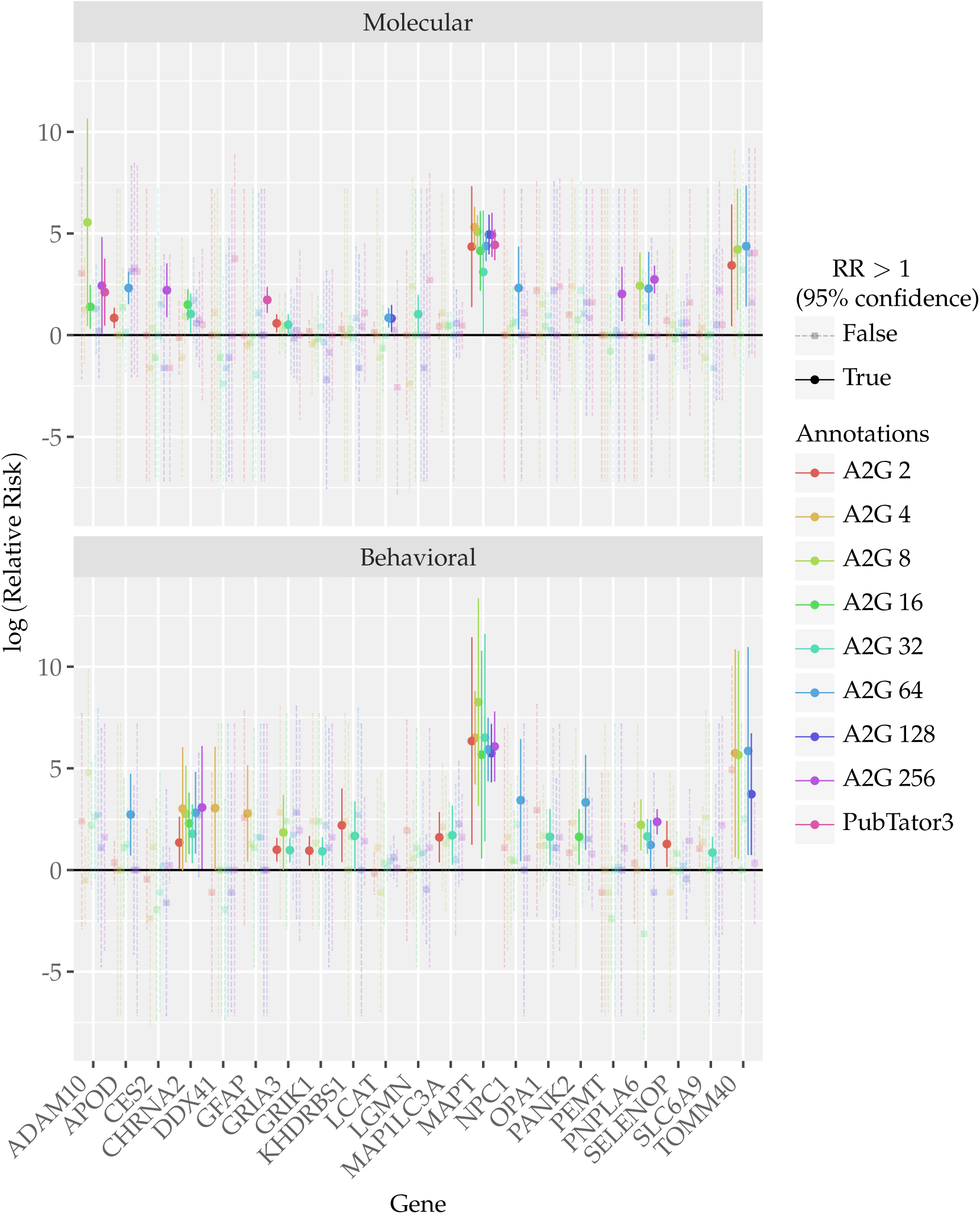
Gene relative risk. Relative risk of a gene being associated with a publication given the publication is related to AD compared to the general publication population. We only display genes where at least one annotation gave a RR significantly greater than 0 (Top) In the molecular test all publications sampled had at least 1 PubTator3 gene annotation. (Bottom) For the behavioral test all publications used were not given any PubTator3 annotations.

We repeated the RR analysis on behavioral publications (Fig 7 bottom). Even in the behavioral case, where PubTator3 did not detect any genes, the models were still able to identify many of the same genes as having a high RR. Notably, *MAPT* was still clearly distinguished by all models. The A2G model trained on 64 labels per batch found the most high-RR genes with 7, 5 of which were the same genes detected in the molecular analysis. These results demonstrate the model’s ability to generalize to abstracts that do not explicitly discuss genes.

## Discussion

In this work, we established A2G an LLM based framework to generate weighted gene predictions for publications which can be used to identify similar abstracts. By taking advantage of the Specter2 LLM’s previously learned ability to compress information from human language into an internal representation, and further training the model to improve its ability to distinguish abstracts based on gene annotations, the model was able to score genes associated with medical abstracts. While we demonstrated value in the model’s predictions, we have identified points that need further investigation as we try to continue progress on this initial release. In the remainder of this discussion, we mention the possibility of a second model for standalone gene annotations and the use of larger LLMs, we question the success of the fine-tuning on masked and unmasked data, and we consider the impact of using different labels per batch during training.

While gene predictions for calculating similarity and creating standalone gene annotations for publications initially seem equivalent, there are some important differences that require unique training strategies to maximize the success of both. In this work, our goal was to find similar publications through gene predictions, but changes to some key decisions in fine-tuning the embedding model would allow new training of the same model architecture to be reused for labeling abstracts. A specific difference between the goal of the search engine and creating standalone annotations that necessitates alternate training is the search engine should bring together related ideas even when the exact genes associated with the publications are not the same, so if we strictly punish the model for giving high scores to genes that are not in a publication but are related to the genes in the publication, we improve gene annotations but hinder the search engines ability to find innovative links. Even without making annotations our primary focus, however, we were still able to get predictions that matched well with the original PubTator3 annotations but we believe this could be improved by creating a second model that focused primarily on predicting gene annotations.

During testing the model it became evident that orthologs and gene families would be a branch point between a model with the goal of predicting gene annotations and another intended for use in a search engine. The decision to convert non-human genes to human orthologs was to improve the search engine but it prevents the models from being used to annotate publications as it’s no longer possible to distinguish between the genes of different species. Without converting to human genes, the model had a hard time differentiating between orthologs and would give similar predictions for multiple species’ genes. For example in AD publications, in addition to the human *APP*, *PSEN1*, *APOE*, etc. the model would make similar predictions for multiple animal *App*, *Psen1*, and *Apoe* genes. When predicting genes for the publication, *Anatomical and functional phenotyping of mice models of Alzheimer’s disease by MR microscopy.* (PMID 17413006) the following predictions greater than 0.1 were given: *Psen1* (0.29), *PSEN1* (0.29), *App* (0.29), *Prnp* (0.29), *BACE1* (0.29), *App* (0.29), *APP* (0.28), *MAPT* (0.26), *APOE* (0.24), *PRNP* (0.24), *Ager* (0.24), *Ldlr* (0.19), *Apoe* (0.16), *Chrna7* (0.14), *Clock* (0.14), *Cnr1* (0.11), *Bmal1* (0.11). There are three variants (including human) of the *APP* gene in the resulting scores, all with the same prediction as well as two *PSEN1* variants with the same predictions and two *APOE* variants with differing predictions. This behavior is consistent, which suggests research on model animals is correlated with analogous human research. It also indicates the model’s training is consistent as it independently predicts similar values for genes across species. For the search engine, this causes issues due to giving extra weight to genes with more orthologs. Secondly, since the search engine should make connections between related work, we want it to connect to publications researching the same topic on different model species. An alternative solution to using orthologs for creating an annotation focused model would be to try to use contrastive learning by creating hard negatives between pairs of publications labeled with orthologs.

Similar to orthologs, we found the issue of gene families, such as the aquaporins (*AQP1*, *AQP2*, *AQP3*, etc). Publication PMID 31624651 (*Distribution of Aquaporins 1 and 4 in the Central Nervous System*) is on *AQP1* and *AQP4* but the model gives high predictions for aquaporins 2, 3, 5, 7, and 9 in addition to 1 and 4. As with orthologs, we could use contrastive learning and select hard negatives by looking for gene symbols that differ only by a number. We can also look at highly co-cited genes or even pull from biological networks like protein-protein interaction networks and gene co-expression networks to find hard negatives to help the model differentiate between genes that have different functions but are often associated together.

To improve the model regardless of goal, a larger embedding model could be used. We chose an embedding model from a set of smaller, ∼100 million parameter, models, which has the obvious advantage of reducing computation cost of both training and embedding all the ∼20 million publications during search engine initiation, but it also provides us with models like SciBERT (and by extension Specter2) and PubMedNCL that were fully pretrained on scientific literature and compiled their own scientific vocabularies. Based on previous research, using a domain-specific vocabulary could be responsible for some of the accuracy improvement over models with vocabularies built from general text [10, 11]. Additional research found pretraining on domain-specific data (including medical literature) could benefit performance compared to starting with weights pretrained on general data [11]. Our own experiments on fine-tuning showed less conclusive results, the unmasked fine-tuning in Fig 9 rank the general purpose models ERNIE 2.0 and BERT as two of the top performing models. This indicates the previous findings may not hold for our use case and using a larger general purpose model may provide easy gains in performance.

When fine-tuning, our goal was to teach the embedding model to ignore gene symbols in text in an attempt to help the model generalize to abstracts without annotations. If we succeeded in this goal, both panels in Fig 9 should look the same, yet for every model except MPNet performance was better on unmasked data. Using the citation network we were able to label some behavioral publications, however, this method was limited to finding enough annotations to bring the annotated data to 5–6% behavioral publications. Reducing standards for annotating behavioral publications or enriching the dataset with behavioral publications from outside the fine-tuning training set would reduce the importance of ignoring gene annotations. Regardless, future work needs to be spent on alternative methods of ensuring the model is capable of handling behavioral publications.

Training the final linear layer with a restricted number of labels was a breakthrough to get non-zero gene predictions at the cost of adding another degree of freedom. We found larger labels per batch reduced predictions globally, leaving only the most confident predictions. Intuitively as the labels per batch increases, a bias for small predictions develops since the sparsity of the labels will lead to zeros dominating the true values. A second reason predictions may be larger for smaller labels per batch is the model is less likely to be punished for predicting an unrelated gene highly. With so many genes, the set of weights that produce a high score for one gene may also cause a related genes to be given a high score. If the number of related genes is low and the number of labels per batch is low, the probability one of these related genes is in the same batch is also low, making it less likely the model will be penalized for the high prediction of a gene not in the publication. Whether this is good or bad depends on the use case of the model. Smaller labels per batch could be seen as giving the model more freedom to stray from the original labels, finding more creative patterns in the relationships between genes. This may help find less obvious connections. Too much freedom, however, and the connections could become too outlandish to be of value. If the goal is to use the model to predict the exact genes related to abstracts, rather than to use the model in the search engine, a larger label per batch would be desirable. It’s also possible varying the labels per batch throughout training could benefit the model but this was not attempted.

Even with these concerns, we exhibited the current models’ ability to make useful gene annotation predictions by comparing the model’s predictions to PubTator3’s gene annotations where the model consistently scored genes higher when the publication had a PubTator3 annotation for that gene compared to publications without the annotation. We showed the model is also able to predict gene annotations in publications without PubTator3 annotations by first running an experiment to compare our predictions on publications to the predictions for papers referencing them. Through a second citation network experiment we found a high correlation between a publication’s top scoring gene prediction and the model’s predictions for the same gene in its references. The ability to predict genes in behavioral publications was further established by our experiment comparing the RR of genes in AD publications calculated using only molecular and then only behavioral publications. We found overlap in genes with RR significantly greater than 0 between the molecular and behavioral cases and the models found more genes with significant risk than the PubTator3 gene annotations. The findings in this experiments provide confidence that the model has learned to make meaningful predictions that can help us search for related research.

## Methods

### Dataset

We based the dataset used throughout this work from the PubTator3 BioCXML archive files. We collected title, PMID, publication year, abstract, and annotations from the full archives. To construct the dataset, we downloaded all 10 archives files from the PubTator3 FTP server (https://ftp.ncbi.nlm.nih.gov/pub/lu/PubTator3). In the BioCXML files some abstracts were split into parts, we concatenated these together to ensure every publication has exactly one abstract.

When collecting annotations, we only used data from the abstracts, excluding those from the title and full text. While our focus was on genes, we wanted to create a dataset that made all the annotations available and make it easy to selectively access that data as needed. PubTator3 identifies several types of annotations, of these types, we tracked gene, disease, species, chemical, cell line, and chromosome annotations. We made a list of identifiers for each annotation type per abstract. In addition to the identifiers, we made a single list of metadata for each abstract. Elements in the metadata list consist of the annotation type, the position, and the length of the annotation in the text, enabling us to find the annotation for corruption. We stored this data in a Hugging Face (HF) dataset and made it available for public use (https://huggingface.co/datasets/dconnell/pubtator3_abstracts) [12].

## Dataset mutation

### Reference

We retrieved partial reference data from PubMed’s FTP server (https://ftp.ncbi.nih.gov/pubmed) using the pubmedparser (https://github.com/net-synergy/pubmedparser) Python package. PubMed’s dataset contained references for only a subset of all publications.

### Gene orthologs

We standardized genes by converting non-human gene IDs to their human orthologs. Using the gene ortholog table available on PubMed’s FTP server, we identified non-human genes and mapped them to human orthologs. Genes that were not in the table were dropped from the annotations.

### Masking

To prevent the model from using gene symbols to make its predictions, we masked the annotations during training by replacing these symbols with ”[MASK]”, using the dataset’s metadata to locate the annotations. We used the same BERT mask symbols for all models despite MPNet using a different string after fine-tuning tests showed MPNet performed comparably with them.

### Permuting

As an alternative form of corruption, in some tests, we permuted annotations by replacing them with random annotations in the dataset. During training time abstract corruption, when permutation was used, we set the permute probability to 0.25 so that 75% of annotations were masked using the mask symbol and the remaining 25% of annotations were permuted. To permute an annotation, we selected a random publication from the dataset and from its annotation list, we picked an annotation of the same type to switch with the target annotation.

### Augmenting behavioral gene annotations

Because, by our definition, behavioral publications lack PubTator3 gene annotations, we inferred annotations for publications heavily referenced by molecular publications, so we could show the model examples of behavioral publications during training. We first filtered the dataset to publications with both PubTator3 annotations and references. Then used the reference data to create a citation graph from these molecular publications to their behavioral references. We then calculated the frequency of each annotation in the molecular publications set to use as the sample probabilities for binomial distributions. For each behavioral publication, we used these binomial distributions to determine the number of references we would expect it to get for each gene at random. If the probability of being cited by at least that many citations with a given gene was below the threshold 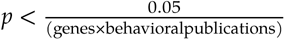 we gave a gene annotation to that publication.

### Maximum number of labels

We excluded publications with > 10 gene annotations under the concern that any publication with that many annotations could not be specific to the genes and would therefore add noise to the models.

### Citation analysis

For the citation analysis, we used the full dataset to determine the distributions of citations between publication types (behavioral or molecular). The PubMed data did not provide reference data for all publications, of 24,593,880 total publications in the dataset, 9,021,257 (or 36.68%) had reference data. It is not clear what factors determine whether reference data exists for a given publication but the proportion of all publications with gene data (27%) is close but less than the proportion of publications with reference data that have gene annotations (31%), suggesting there is a bias towards molecular publications. This discrepancy could have impacted the final results of this analysis.

### Templates

We create the templates by passing 32 examples of abstracts with the PubTator3 gene annotation through the embedding unit and averaging their embeddings (Fig 1 C). We make predictions by comparing the test abstract’s embedding to each gene template using a dot product then use a sigmoid function to transform the results into a probability in the range (0, 1).

While a multi-label Neural Network (NN) could directly predict gene annotations without templates, we found several reasons to use templates. Templates stiffen the model reducing the risk of overtraining. Neural networks are flexible and since we are concerned the PubTator3 annotations may be noisy, training a flexible model risks instilling patterns of incorrect labels in the final weights. A stiffer model limits the model to learning only the strongest signals. Secondly, by using templates, every training abstract modifies the same weights. This improves the model’s ability to make predictions on genes with few publications by reusing the information gained from other publications.

### Running experiments

To ensure experiments are not impacted by using training data, the dataset is broken into multiple single use splits. We split data into separate splits for training the embedding model, training the A2G model, the label similarity analysis, comparing against PubTator3 annotations, and the DE experiment. The reference analysis is an exception since we need to collect all references from publications and these are scattered across the dataset.

To make the research reproducible, we assign a random seed to each script. We started with a random seed of 10 for the first script then all sequential scripts get a seed of 10 more than the previous script to allow scripts to use up to 10 seeds for multiple random number generals. In a few scripts when several random number generators were used, we created a seed generator with the script’s seed then used this to derive seeds for other random number generators to simplify managing multiple seeds.

### Embedding model fine-tuning and selection

Before training the A2G model, we experimented with approaches to fine-tune and select between possible embedding models. Base models are trained for general document representation. To maximize performance on distinguishing abstracts by gene, we can ”fine-tune” the model by continuing training on a specific task. Our goal was to create a task that would maximize similarity between abstracts with a common gene and minimize similarity between abstracts without a common gene. We ran experiments on different fine-tuning strategies for 7 embedding models to find the best model and training method.

The embedding model converts variable-length abstracts to a fixed-width numerical vector that can be used with traditional machine learning approaches. To find the best model, we started by selecting hyperparameters, used those hyperparameters to do an initial set of fine-tuning method experiments on two of the models, then trained each model using the selected fine-tuning method.

The seven embedding models were split between general purpose models (those pre-trained on generic text) and scientific models (trained on scientific literature). We started with the two top performing general purpose embedding models in a test performed by the sentence-bert maintainers [13], Microsoft’s MPNet (mpnet-base) [14] and PaddlePaddel’s ERNIE 2.0 (ernie-2.0-en) [15] and added Google’s BERT (bert-base-uncased) [16]. For the scientific models, we added four more models based on BERT’s architecture: Allen Institute for AI’s SciBERT uncased (scibert_scivocab_uncased) [10], Specter [17], and newer Specter2 [8], and PubMedNCL [18] which is a fine-tune of Microsoft’s BiomedBert—a model similar to SciBERT but pretrained from PubMed data instead of the Semantic Scholar dataset. In addition to being trained on scientific literature, the tokenizers’ vocabularies were also generated from domain-specific text, which improves performance [10].

### Hyperparameter selection

We selected hyperparameters through a series of 30 short fine-tune trials for each model with values selected at random [19]. From the trials we chose values for the following hyperparameters: batch size, warm-up ratio, and learning rate. Each parameter was given a specified search space that values could be sampled from. For batch size, the search space was integers between 8 and 64. Warm-up ratio’s search space was floating precision values between 0 and 0.3 and for learning rate, the search space was between 1 × 10^−6^ and 1 × 10^−4^. We used the hyperparameters from the best performing trial for the remainder of the experiments. The final hyperparameters used are presented in Table 2.

**Table 2.** The hyperparameters that resulted in the best performance across 30 short fine-tuning trials.

| Model | Batch Size | Warmup Ratio | Learning Rate |
| --- | --- | --- | --- |
| ERNIE 2.0 | 44 | 0.0480 | $3.918 \times 10^{-5}$ |
| MPNet | 58 | 0.2109 | $5.653 \times 10^{-5}$ |
| BERT | 32 | 0.0519 | $4.875 \times 10^{-5}$ |
| SPECTER | 42 | 0.1995 | $4.414 \times 10^{-6}$ |
| SPECTER2 | 16 | 0.1198 | $5.459 \times 10^{-6}$ |
| SciBERT | 8 | 0.0665 | $3.222 \times 10^{-6}$ |
| PubMedNCL | 15 | 0.1574 | $1.698 \times 10^{-5}$ |

To perform the trials, we used the optuna Python package [20]. Because we performed the hyperparameter selection before the fine-tuning experiments, we used a dataset with gene annotations masked instead of masking both gene and disease annotations as we used in later fine-tuning.

We used a fixed 64,000 examples for all trials and calculated the number of steps based on the batch sizes to prevent trials with larger batch sizes from seeing more examples. For batches, we created anchor–positive–negative triplets where a randomly selected label was chosen for each triplet, ensuring that a label could only be drawn once per batch, and we sampled two publications, the anchor and the positive, from the pool of publications annotated with that label. We then selected a negative at random ensuring it did not have the label as an annotation. We did not correct for the possibility that a negative selected might have a different label in common with the anchor.

During the triplet selection process, we selected labels using the log of their frequency as weights. Since the most commonly written about genes are orders of magnitude more prevalent in the literature than the least common genes, we did not want a few ”star” genes to dominate the dataset as would be the case if selection used their raw frequency. In contrast, we didn’t want the same few publications to be used whenever a rarely written about gene was selected as the triplet label in the case where a uniform distribution was used for selecting labels at random. So selection based on the log of their frequency was used as a compromise to ensure a good mix of both genes and publications in the dataset.

For the training, we dropped the negative values and used a loss function that maximizes the similarity between each anchor–positive pair while minimizing the similarity between the anchor and the positives of the other pairs in the batch [21]. When evaluating, the accuracy was computed based on whether the anchor and positive were more similar than the anchor and negative for a given triplet.

### Fine-tuning experiments

Before performing the final fine-tunings, we performed a suite of experiments to determine the best task to fine-tune on for our purposes. To improve the model’s ability to generalize to publications without PubTator3 annotations we wanted to prevent the model from depending on gene symbols in the abstracts, leading to masking the symbols. In contrast, the model should perform well on unmasked abstracts since the trained model will be given raw abstracts.

Because of this, it’s not obvious whether fine-tuning should be performed on masked or unmasked data so we devised experiments that track how well embedding models fine-tuned using different strategies performed on masked and unmasked.

We performed 5 experiments on a general purpose model, MPNet, and a scientific model, PubMedNCL, chosen based on their performance on early hyperparameter tests. The five experiments were fine-tuning with 1. no masking (control), 2. genes masked, 3. genes and disease masked, 4. genes and disease masked but alternating the task, and 5. genes and disease corrupted through masking and permutation. We excluded the fourth experiment from the results figure due to poor performance. For each experiment, we evaluated the models on three datasets: no masking, genes masked, and genes and diseases masked. We ran 10,000 training steps and evaluated the model every 250 steps. We used the same methods for creating datasets and the same multiple negative loss function as we used during hyperparameter selection.

We created the last experiment after noting that fine-tuning on masked data hindered the model’s performance on unmasked abstracts. This can be seen in Fig 8, horizontal line 1 shows the performance on unmasked data fell as fine-tuning on masked data continued. Our intention was to allow the model to see abstracts both with and without gene symbols but train it to ignore the gene symbols by ensuring they provide no information.

**Figure 8.**
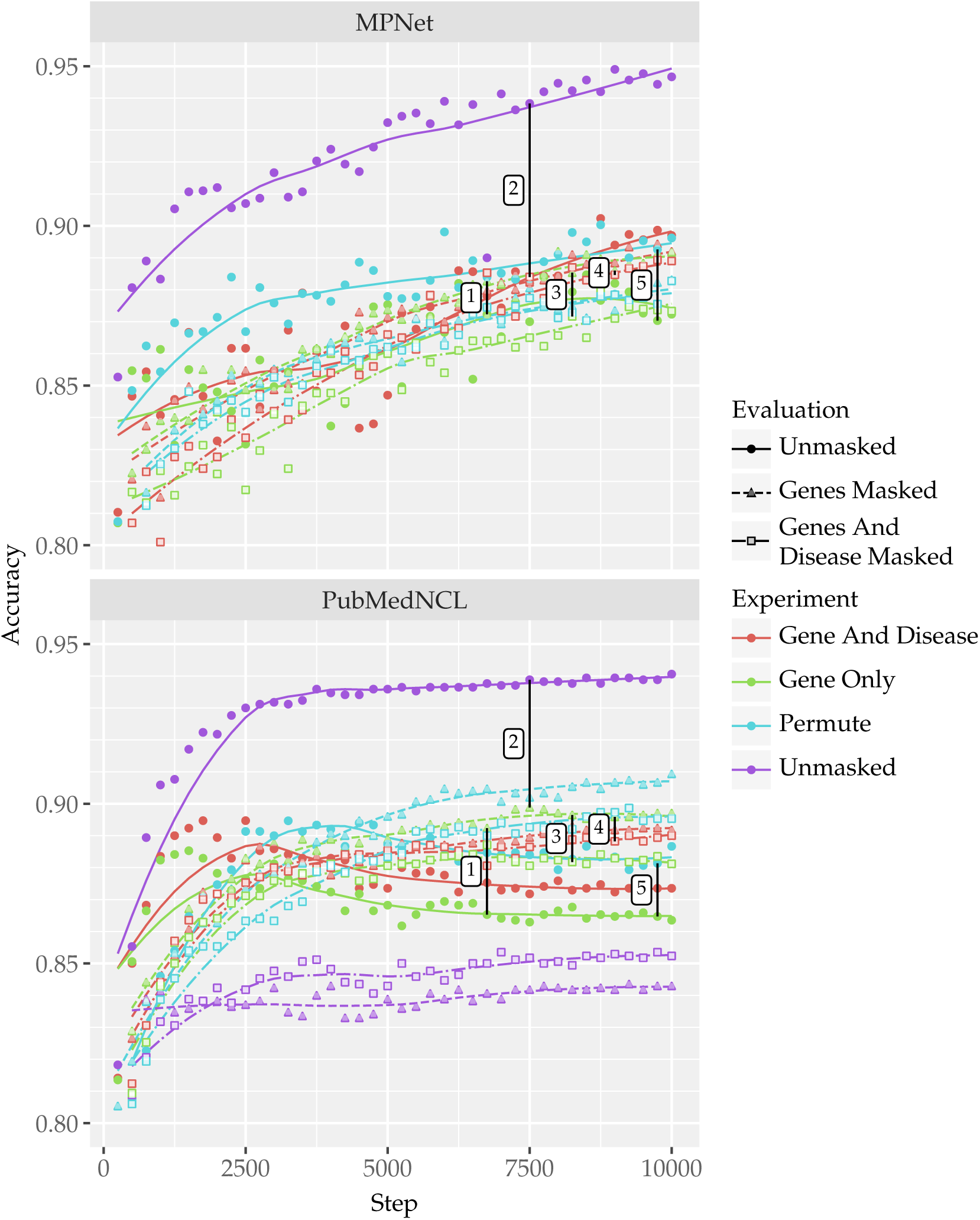
Comparison of fine-tuning strategies. Training curves of MPNet (top) and PubMedNCL (bottom) over 10,000 training steps and four experiments. In both figures, line 1 highlights the difference in the gene only’s performance between evaulating on a gene masked and unmasked dataset. Line 2 is the cost of masking genes. Line 3 is the difference between evaulating the gene only experiment on gene masked dataset vs a gene and disease masked dataset. Line 4 is similar to three but it compares the gene only experiments performance on the gene masked dataset to the gene and disease experiments performance on the gene and disease masked dataset. Line 5 is the improvement permutation during fine-tuning provided to performance on the unmasked dataset.

The results of the experiments are shown in Fig 8. When comparing the training curves of MPNet and PubMedNCL, PubMedNCL improves at a faster rate early on but approaches an asymptote much sooner than MPNet, which is still trending upwards even at the end of the 10,000 training steps.

For both models, performance was poor on masked datasets after fine-tuning on an unmasked dataset (for MPNet the masked evaluations’ accuracies are below the plot area, around 0.7). This highlights the importance of masking the data for generalization since the model is relying on gene symbols in abstracts. As line 1 shows, the converse is also true. We see the same pattern when the model was trained on gene masked data, the performance on unmasked data is worse. This leads to a trade off in performance on behavioral vs molecular abstracts. The MPNet model has a much smaller difference in performance between predictions on masked and unmasked data when trained on gene masked (or gene and disease masked) data. This could be due to PubMedNCL’s prior knowledge of genes causing the sight of the genes to influence the final embedding more significantly while MPNet gives low attention to gene names having seen few examples of them. While it is difficult to see, looking closely at the unmasked evaluation lines, from start to around 2500 steps, PubMedNCL’s performance on the unmasked dataset when trained with masked genes increases, peaks, then starts to decline with further training. So additional fine-tuning reduces PubMedNCL’s ability to make predictions on unmasked data, a pattern not seen in MPNet. In each of the four experiments trained on masked data, the unmasked evaluation lines are distinctly less than the masked. For MPNet these lines are mixed.

Line 2 compares the performance of a model fine-tuned on gene masked data to a model fine-tuned on unmasked data when evaluated on the type of data they are fine-tuned on. This shows both embedding models can be trained to pick out gene names and use them to influence the final embedding. This line confirms that training on unmasked data would inhibit generalization to gene-free abstracts since the model relies heavily on the presence of gene names in creating embeddings. At the first evaluation point (step 250), MPNet’s unmasked performance is ahead of PubMedNCL. After that its training is slower but it passes PubMedNCL again after PubMedNCL’s curve flattens out.

Line 3 is the difference between the performance on gene masked data and gene and disease masked data by the models trained on gene masked data. This is intended to answer whether the gene masked model uses disease names as a proxy for gene names. In both models, the difference between the two evaluation methods is similar. Performance is improved by the presence of disease names so it will likely be beneficial for the search engine purposes to remove diseases from the training dataset for the same reasons genes need to be redacted. Promisingly, line 4 shows, when trained without disease names, the performance on abstracts with gene and disease masked gets closer to the performance on abstracts with only genes removed. In MPNet, this difference is particularly small.

Line 5 indicates how well the permutation fine-tuning strategy reduced the damage fine-tuning on masked data has on the model’s ability to predict unmasked data. For both embedding models, the unmasked performance raises compared to the standard gene masked training. There is, however, the same trend in PubMedNCL of the unmasked performance peaking earlier in fine-tuning then falling. For MPNet permutation training does improve the performance on unmasked data without hurting the performance on masked data such that the gene masked and unmasked data are about the same as opposed to the standard masked training lines where performance on masked data was a little better.

These experiments suggest we should train on gene and disease masked data since removing diseases from the abstracts does not have much impact on performance and could help improve generalization. Additionally, MPNet does not appear to have hit a plateau by 10,000 training steps and could benefit from further fine-tuning. MPNet is not without downsides however. It is much slower, fine-tuning took about 2.5 times as long as the other models and as Fig 8 shows, the performance has more variance than PubMedNCL.

### Model selection

Based on the experiments, we found the masking and permutation method of corruption to be the best approach to training, with corruption performed on both genes and diseases. We fine-tuned all 7 models using only this approach and 20,000 steps to make sure the training curves flatten out. Fig 9 shows the results of this final fine-tuning experiment.

**Figure 9.**
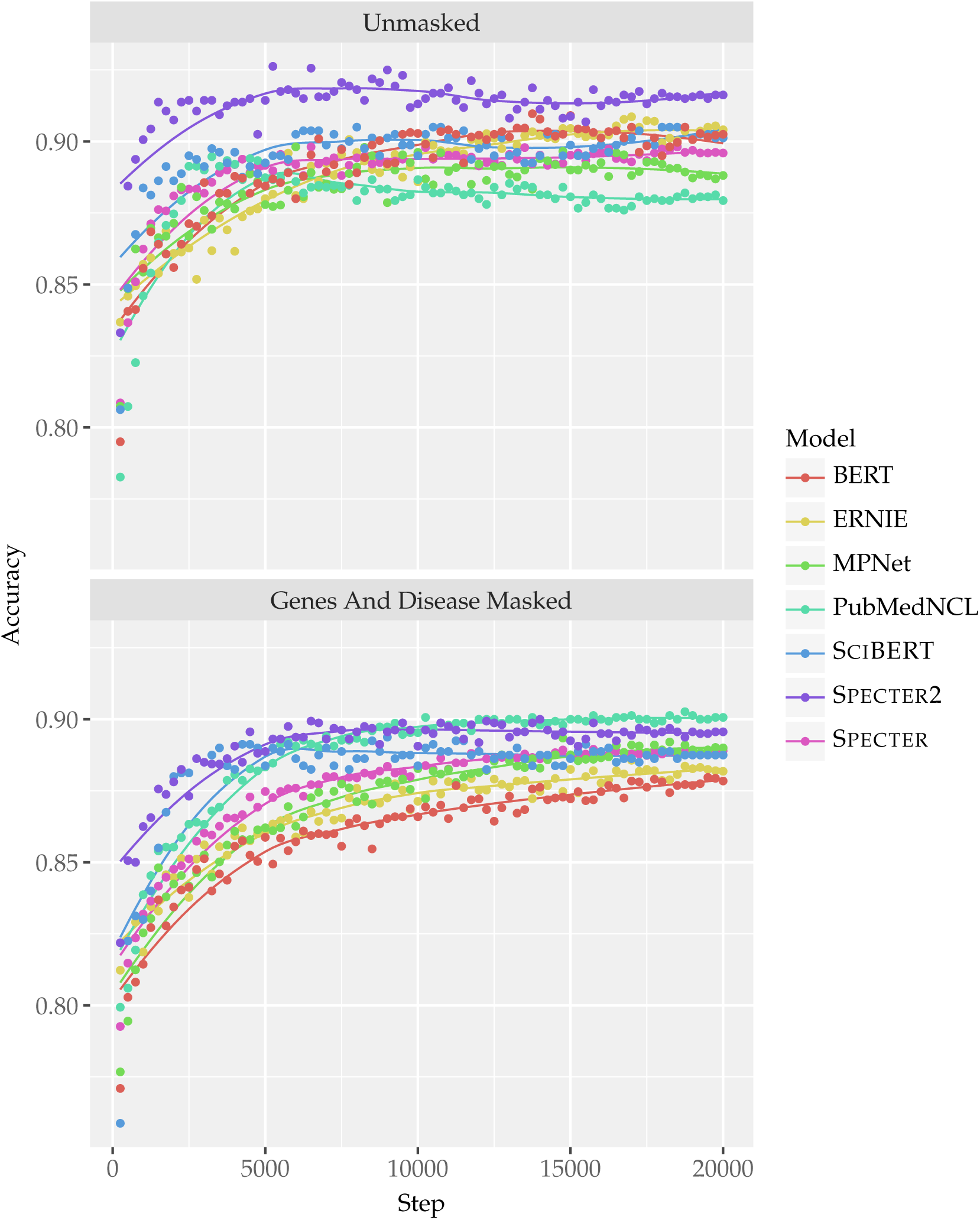
Training curve comparison. Results of the fine-tuning each of the test models evaluated on unmasked data (top) and gene and disease masked data (bottom). Fine-tuning was performed with gene and disease masked data using the permute training approach.

In Fig 9, the unmasked evaluation represents the model’s quality on molecular studies while the masked is an attempt to represent how the model performs on behavioral references in lieu of annotations. Specter2 performs best in the unmasked case and is a close second in the masked case leading us to choose this model for A2G. Notably, the relationship between performance on unmasked and masked data varies between the models. PubMedNCL performs best on masked data but worst on unmasked, yet another scientific model, Specter2, performs better on the unmasked data than it does on the masked data. Surprisingly, in several cases general purpose models outperform the domain specific models, despite the scientific models having an advantage of a domain-specific vocabulary.

### Label embedding similarity

We tested the impact of fine-tuning the Specter2 model by comparing the original (”base”) and fine-tuned models’ ability to produce similar embeddings for abstracts labeled with a common gene. For this experiment, we selected 15 genes with at least 200 example publications at random. Pooling all publications from the 15 genes, we discarded any annotated with more than 1 of the chosen genes before sampling 200 publications for each gene. We passed all the 15 × 200 abstracts through both embedding models and averaged embeddings in groups of four to simulate templates, resulting in 50 average embeddings for each gene. To score similarity, we used Pairwise correlation on all embeddings.

Fig 10, compares the distribution of correlations between embeddings from the same gene and embeddings with different genes. The base model created embeddings that had a higher overall similarity. This may be due to the model being trained on a large corpus of scientific literature leading it to view molecular publications as similar within the more general set. Due to the higher average similarity, we normalized the correlation matrices by removing the mean and dividing by the standard deviation. Before and after fine-tuning the similarity between embeddings for publications given the same PubTator3 annotation tended to be higher than those without a common annotation. After fine-tuning, the distribution of similarities in the common gene case is moved higher while the distribution of similarities among publications without a common gene is moved down.

**Figure 10.**
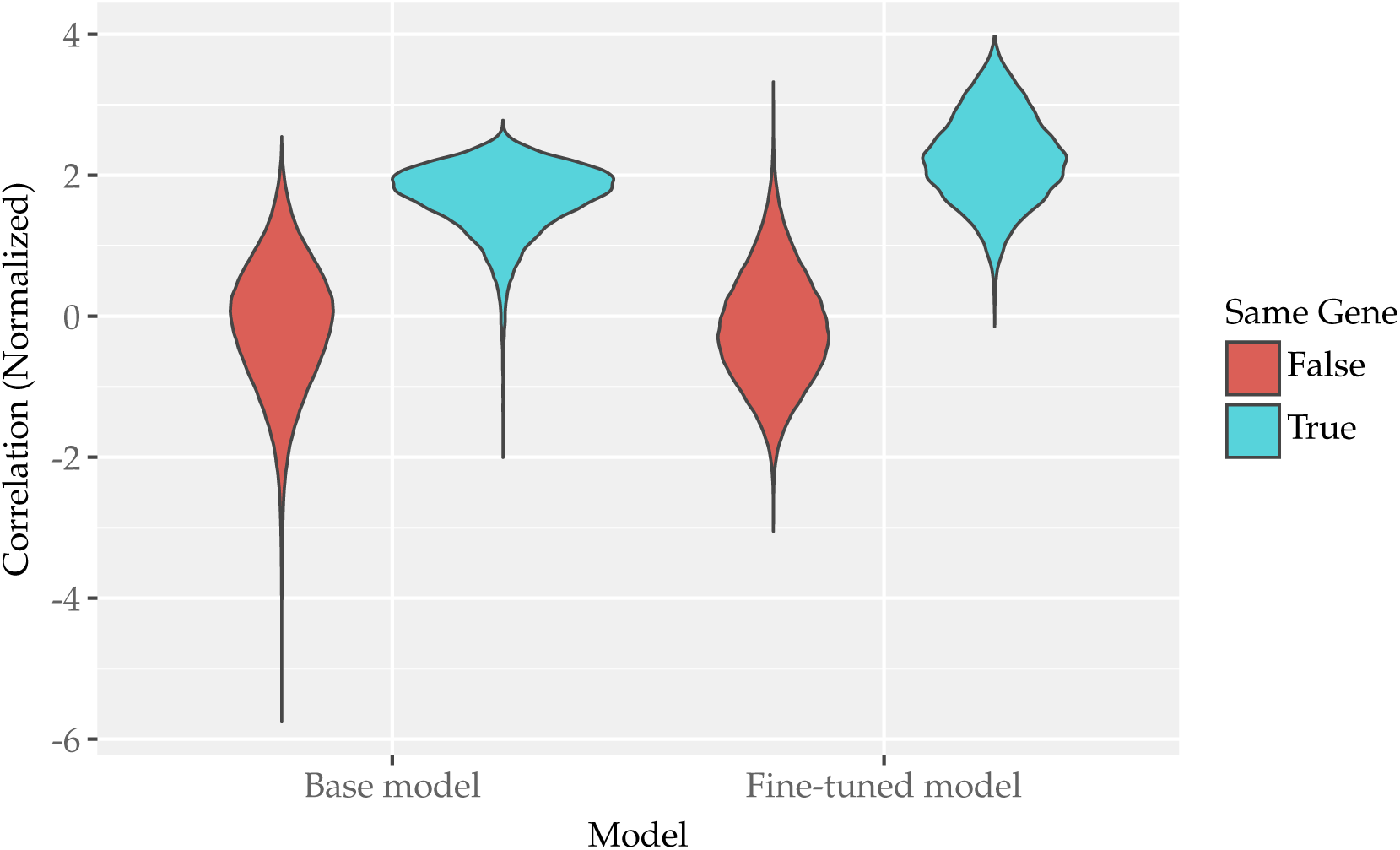
Label similarity before and after fine-tuning. Normalized cosine similarties of publication embeddigs created with the Specter2 model before and after task specific fine-tuning. Distributions are created for similarity between publication pairs that share a gene annotation and those without a common gene.

### A2G training

The A2G model is composed of a 768 node linear layer on top of the embedding model. This combination of embedding model, linear layer and GeLU gate comprises the ”embedding unit” which returns a numeric vector of length 768. Both the gene template abstracts and input abstracts are sent through this embedding unit before being compared by a dot product resulting in one value for each gene template. After taking the dot product with the gene templates, a trained scalar bias is added to the results which then go through a sigmoid function to transform the results to a value between 0 and 1.

We used jax (https://github.com/jax-ml/jax) for training along with the higher level packages built on-top of jax, flax and optax. For updating weights, we used the ADAM optimizer with a learning rate of 1 × 10^−4^ and a sigmoid binary cross entropy loss function. During training, we made batches of input abstracts, template abstracts, and input labels. We created batches during training with the labels for the current batch selected at random from the pool of all labels that have not yet been used during the current epoch. Epochs lasted until the batches exhausted the available labels.

The input labels were a boolean vector of length labels per batch with 1 meaning the input abstract was annotated with the gene at that index. While we selected input abstracts for a single label in the batch, it’s possible an abstract could have annotations for multiple labels in the batch (i.e. more than one non-zero value in the input labels).

To fully train the model, we performed multiple training loops with varying template sizes. We stopped a training loop when either a specified number of epochs occurred or if the rolling average change in the test loss between epochs was less than 1 × 10^−5^. We started training with a high noise template size of 1 for 20 epochs then we increased the template size to 4, 8, and then the final template size of 32 for 10 epochs each.

Once training was finished, we ”attached” the templates to the model by sampling template size number of examples per gene in the dataset and sending them through the embedding unit then averaging the results, since the weights of the linear layer are now fixed and the template vectors no longer change, after training these can be stored with the model and don’t need to be recomputed. The dataset split we used during training is non-overlapping with the dataset splits used in the model testing experiments (except in the behavioral publication gene prediction experiment where the full dataset must be used to get the references) which prevents the chance an abstract used in the experiments was used to create the templates. We trained models using 2, 4, 8, 16, 32, 64, 128, and 256 labels per batch to determine how labels per batch impacts model behavior.

### Comparison to PubTator3

We used the models to perform predictions on 30 examples each of abstracts annotated with 18 randomly selected genes and an equal number of random publications. For each gene, we then compared the score for the 30 gene annotated publications to the 30 random publications. For the model comparison, we reused the same data but aggregated it over the model so each model had 30 × 18 examples of matches and the same number of non-matching data.

### Predicting genes in behavioral publications

To determine if the A2G model can detect meaningful gene components from abstracts that PubTator3 did not find gene annotations for, we designed an experiment to compare the gene predictions of a molecular ”parent publication” to the gene predictions of its references. From the dataset, we selected 10,000 parent publications and filtered to publications fitting the requirements of having at least one PubTator3 annotation and having reference data. This resulted in 1120 parent publications. For each parent publication, we selected a single behavioral reference at random along with a molecular reference and a random unrelated behavioral publication to use as benchmarks. We then sent all selected publications through each A2G model to predict their genes. Using the resulting predictions, we correlated the child publications’ gene vectors with their parent publications’ predictions using Pearson correlation.

We performed a second analysis to determine if the prediction for the parent publication’s top scoring gene was correlated with the behavioral references prediction for the same gene. To do this, we determined the top scoring gene for each parent publication and found the corresponding prediction for each of its three child publications then plotted the scores of the children against their parents score.

### Alzheimer’s Disease Differentially expressed genes

To calculate DE genes, we obtained bulk transcriptome data from the Religious Order Study and Memory and Aging Project (ROSMAP) study collected from the medial frontal cortex of study participants along with labels for their cognitive state [9]. Cognitive state scores range from 1 (NCI) to 4 (AD). We split data into NCI and AD, discarding data for cognitive states of 2 and 3. Using a two-sided t-test, we then compared the gene expression between both groups using an 𝛼 of 0.05 and Bonferroni correction to handle multiple tests, resulting in 296 potential genes with differential expression associated with AD.

We separated publications into four groups based on two criteria, AD vs non-AD and molecular vs behavioral. To identify AD related publications, we searched for the term ”Alzheimer” in the paper’s abstract. This resulted in 10,738 molecular and 8799 behavioral AD abstracts. We sampled an equal number of molecular and behavioral non-AD respectively.

For molecular publications, we collected PubTator3 annotations and calculated the proportion of gene annotations as the sum of annotations divided by the number of DE genes × the number of publications. For each model, we selected a threshold using the number of PubTator3 annotations as a target. We then passed the title and abstract of all publications to each A2G model and the calculated threshold was applied to the predictions to determine the number of events (genes predicted) and non-events for each gene and annotation method.

We needed the threshold to make direct comparisons to the PubTator3 annotations and to calculate RR. After binarizing the data, we estimated the RR for each gene associated with AD. We chose those that had an RR > 1 with 95% confidence in any of the annotation methods to to be displayed in Fig 7. We use the equation for SE (log(RR)) to calculate the Confidence Interval (CI) (Eq. (1)).

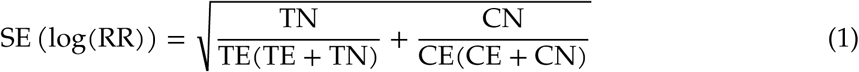

Where log is the natural log, TN is the treatment non-events, TE treatment events, CN control non-events, and CE control events. Inside the square root of Eq. (1) there are terms for the treatment (AD) group and the control group. The two terms can be split into the proportion of all samples in a group that are a non-event (in the control case 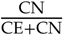) and the inverse of the absolute number of samples that are an event, 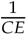. There are then two ways to reduce the standard error, reduce the proportion of non-events (increase the proportion of events) and increase the total number of events. Increasing the total number of events is the intuitive method for reducing the Standard Error (SE), under a constant proportion of events this means increasing the sample size. The other method, increasing the proportion of events, means a model that predicts genes more frequently will consequently have a lower SE. (Additionally, increasing the proportion of events will lead to a smaller 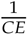 as well, further reduces SE.) Assuming the population RR is the same, the statistics will favor a model that predicts more genes. To correct for this, we calculate the proportion of all possible annotations PubTator3 predicts (approximately 8 × 10^−4^) and then for each model determine the threshold that will produce the same prediction proportion in that model. For A2G models trained with 2, 4, 8 labels per batch, the threshold is > 0.99.

After adding 0.5 to both the event and non-event counts to prevent divide-by-zero errors, we used the events and non-events to calculate the RR for each gene and the SE for the gene. We performed a one sided t-test corrected for multiple tests to find the genes that had a log RR significantly greater (𝛼 = 0.05) than 0.

### Search engine

We built the search engine with fastapi (https://github.com/fastapi/fastapi) to create the web application and qdrant (https://github.com/qdrant/qdrant) as a vector database to cache predictions. Based on a combination of the experiments and testing on subsets of the full dataset, we chose the 64 label per batch model for use in the search engine. On the first run, we populated the qdrant database by using the A2G model to make predictions for all the 24 million publications in the PubTator3 archive dataset and then sent those vectors to the database along with publication title, year, the raw abstract, PubTator3 gene annotations, and reference data.

Once running, a user can submit a new title and abstract to the web UI and the search engine will predict genes with A2G model and pass the resulting vector to qdrant’s search function to get a list publication IDs whose gene predictions most closely match the user’s input based on cosine similarity.

## Data availability

The PubTator3 dataset created in this work can be found at https://huggingface.co/datasets/dconnell/pubtator3_abstracts. The fine-tuned Specter embedding model is also available on HF, https://huggingface.co/dconnell/SPECTER2-abstract2gene

## Code availability

The code used to fine-tune the models, perform the experiments, and run the web UI can be found at https://github.com/net-synergy/abstract2gene.

## Supplementary Information

**Figure S1.**
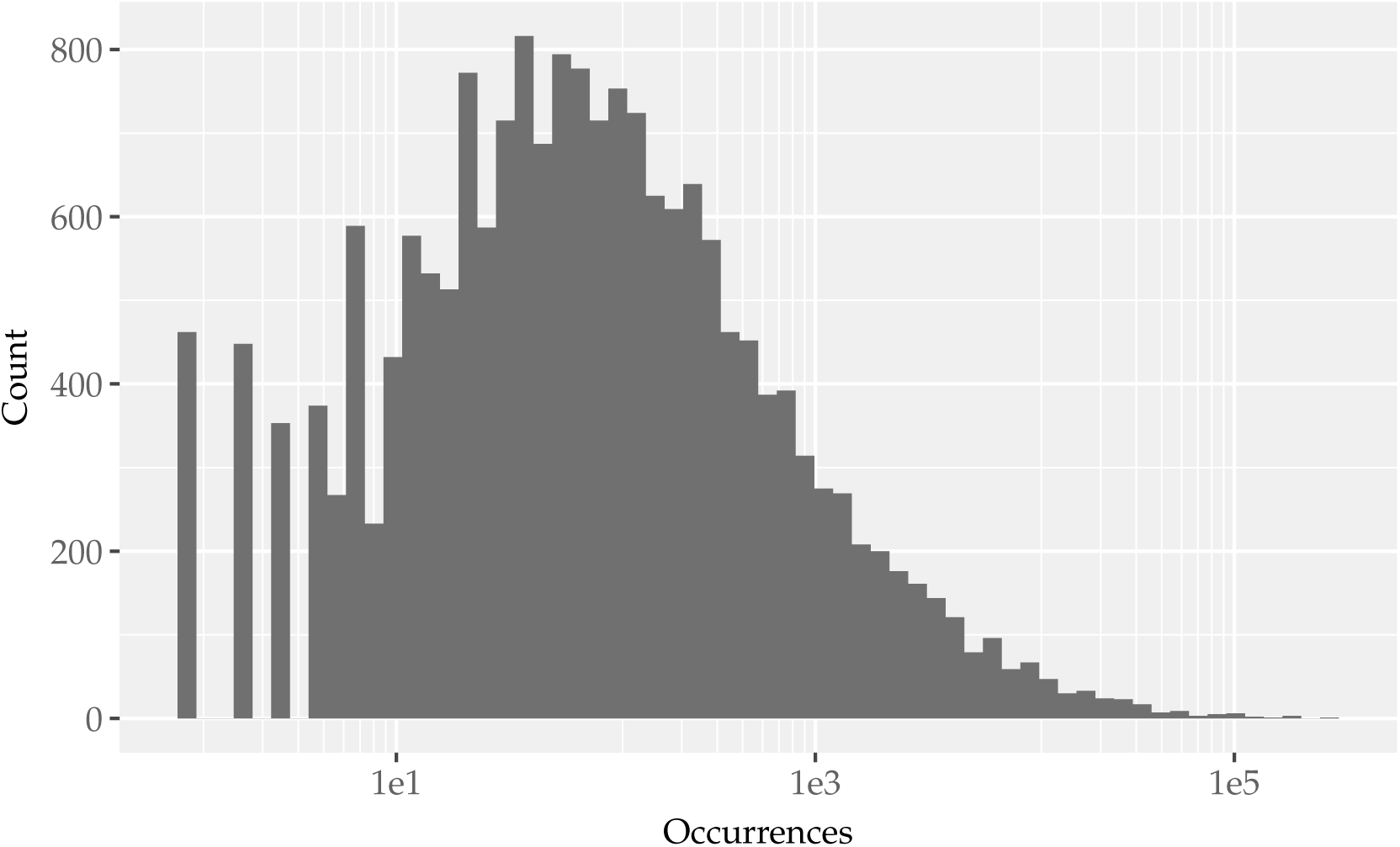
Distribution of gene annotation prevalance in the PubTator3 abstract dataset.

## References

1. Zhou X, Menche J, Barabási AL, Sharma A. Human Symptoms-Disease Network. Nature Communications. 2014;5(1):4212. doi:10.1038/ncomms5212.

2. Barrenas F, Chavali S, Holme P, Mobini R, Benson M. Network Properties of Complex Human Disease Genes Identified Through Genome-Wide Association Studies. PLoS ONE. 2009;4(11):e8090. doi:10.1371/journal.pone.0008090.

3. Goh KI, Cusick ME, Valle D, Childs B, Vidal M, Barabási AL. The Human Disease Network. Proceedings of the National Academy of Sciences. 2007;104(21):8685–8690. doi:10.1073/pnas.0701361104.

4. Jeong H, Mason SP, Barabási A, Oltvai ZN. Lethality and Centrality in Protein Networks. Nature. 2001;411(6833):41–42. doi:10.1038/35075138.

5. Cheng D, Xue Z, Zhibo Y, Mingze Z. Impact of Interdisciplinarity on Disruptive Innovation: the Moderating Role of Collaboration Pattern and Collaboration Size. Scientometrics. 2025;130(4):2379–2401. doi:10.1007/s11192-025-05285-3.

6. Rzhetsky A, Foster JG, Foster IT, Evans JA. Choosing Experiments To Accelerate Collective Discovery. Proceedings of the National Academy of Sciences. 2015;112(47):14569–14574. doi:10.1073/pnas.1509757112.

7. Wei CH, Allot A, Lai PT, Leaman R, Tian S, Luo L, et al. Pubtator 3.0: an Ai-Powered Literature Resource for Unlocking Biomedical Knowledge. Nucleic Acids Research. 2024;52(W1):W540–W546. doi:10.1093/nar/gkae235.

8. Singh A, D’Arcy M, Cohan A, Downey D, Feldman S. Scirepeval: a Multi-Format Benchmark for Scientific Document Representations. arXiv preprint arXiv:221113308. 2022;doi:10.48550/ARXIV.2211.13308.

9. Bennett DA, Buchman AS, Boyle PA, Barnes LL, Wilson RS, Schneider JA. Religious Orders Study and Rush Memory and Aging Project. Journal of Alzheimer’s Disease. 2018;64(s1):S161–S189. doi:10.3233/jad-179939.

10. Beltagy I, Lo K, Cohan A. SciBERT: A Pretrained Language Model for Scientific Text. In: Proceedings of the 2019 Conference on Empirical Methods in Natural Language Processing and the 9th International Joint Conference on Natural Language Processing (EMNLP-IJCNLP); 2019.Available from: 10.18653/v1/D19-1371.

11. Gu Y, Tinn R, Cheng H, Lucas M, Usuyama N, Liu X, et al. Domain-Specific Language Model Pretraining for Biomedical Natural Language Processing. ACM Transactions on Computing for Healthcare. 2021;3(1):1–23. doi:10.1145/3458754.

12. Lhoest Q, Villanova del Moral A, Jernite Y, Thakur A, von Platen P, Patil S, et al. Datasets: A Community Library for Natural Language Processing. In: Proceedings of the 2021 Conference on Empirical Methods in Natural Language Processing: System Demonstrations. Online and Punta Cana, Dominican Republic: Association for Computational Linguistics; 2021. p. 175–184. Available from: https://aclanthology.org/2021.emnlp-demo.21.

13. Reimers N, Gurevych I. Sentence-Bert: Sentence Embeddings Using Siamese Bert-Networks. arXiv preprint arXiv:190810084. 2019;doi:10.48550/ARXIV.1908.10084.

14. Song K, Tan X, Qin T, Lu J, Liu TY. Mpnet: Masked and Permuted Pre-Training for Language Understanding. CoRR. 2020;.

15. Sun Y, Wang S, Li Y, Feng S, Tian H, Wu H, et al. Ernie 2.0: a Continual Pre-Training Framework for Language Understanding. CoRR. 2019;.

16. Devlin J, Chang MW, Lee K, Toutanova K. BERT: Pre-training of Deep Bidirectional Transformers for Language Understanding. In: Proceedings of the 2019 Conference of the North American Chapter of the Association for Computational Linguistics: Human Language Technologies, Volume 1 (Long and Short Papers); 2019. p. 4171–4186. Available from: 10.18653/v1/N19-1423.

17. Cohan A, Feldman S, Beltagy I, Downey D, Weld DS. Specter: Document-Level Representation Learning Using Citation-Informed Transformers. CoRR. 2020;.

18. Ostendorff M, Rethmeier N, Augenstein I, Gipp B, Rehm G. Neighborhood Contrastive Learning for Scientific Document Representations With Citation Embeddings. arXiv preprint arXiv:220206671. 2022;doi:10.48550/ARXIV.2202.06671.

19. Bergstra J, Bengio Y. Random Search for Hyper-Parameter Optimization. J Mach Learn Res. 2012;13:281–305.

20. Akiba T, Sano S, Yanase T, Ohta T, Koyama M. Optuna. In: Proceedings of the 25th ACM SIGKDD International Conference on Knowledge Discovery and Data Mining; 2019. p. 2623–2631. Available from: 10.1145/3292500.3330701.

21. Henderson M, Al-Rfou R, Strope B, Sung Yh, Lukacs L, Guo R, et al. Efficient Natural Language Response Suggestion for Smart Reply. CoRR. 2017;.

